# A genomic region containing *REC8* and *RNF212B* is associated with individual recombination rate variation in a wild population of red deer (*Cervus elaphus*)

**DOI:** 10.1101/248468

**Authors:** Susan E. Johnston, Jisca Huisman, Josephine M. Pemberton

## Abstract

Recombination is a fundamental feature of sexual reproduction, ensuring proper disjunction, preventing mutation accumulation and generating new allelic combinations upon which selection can act. However it is also mutagenic, and breaks up favourable allelic combinations previously built up by selection. Identifying the genetic drivers of recombination rate variation is a key step in understanding the causes and consequences of this variation, how lociassociated with recombination are evolving and how they affect the potential of a population to respond to selection. However, to date, few studies have examined the genetic architecture of recombination rate variation in natural populations. Here, we use pedigree data from ‐2,600 individuals genotyped at ‐38,000 SNPs to investigate the genetic architecture of individual autosomal recombination rate in a wild population of red deer (*Cervus elaphus*). Female red deer exhibited a higher mean and phenotypic variance in autosomal crossover counts (ACC). Animal models fitting genomic relatedness matrices showed that ACC was heritable in females (*h*^2^ = 0.12) but not in males. A regional heritability mapping approach showed that almost all heritable variation in female ACC was explained by a genomic region on deer linkage group 12 containing the candidate loci *REC8* and *RNF212B*, with an additional region on linkage group 32 containing *TOP2B* approaching genome-wide significance. The *REC8/RNF212B* region and its paralogue *RNF212* have been associated with recombination in cattle, mice, humans and sheep. Our findings suggest that mammalian recombination rates have a relatively conserved genetic architecture in both domesticated and wild systems, and provide a foundation for understanding the association between recombination loci and individual fitness within this population.

## Introduction

Meiotic recombination (or crossing-over) is a fundamental feature of sexual reproduction and an important driver of diversity in eukaryotic genomes (Felsenstein, 1974; Barton and Charlesworth, 1998). It has several benefits: it ensures the proper disjunction of homologous chromosomes during meiosis (Hassold and Hunt, 2001), prevents mutation accumulation (Muller, 1964) and generates novel haplotypes, increasing the genetic variance for fitness and increasing the speed and degree to which populations respond to selection (Hill and Robertson, 1966; Battagin *et al*., 2016). However, recombination can also come at a cost: it requires the formation of DNA double strand breaks which increase the risk of local mutation and chromosomal rearrangements (Inoue and Lupski, 2002; Arbeithuber *et al*., 2015); it can also break up favourable allele combinations previously built up by selection, reducing the mean fitness of successive generations (Barton and Charlesworth, 1998). Therefore, as the relative costs and benefits of recombination vary within different selective contexts, it is expected that recombination rates should vary within and between populations (Burt, 2000; Otto and Lenormand, 2002). Indeed, recent studies have shown that recombination rates can vary within and between chromosomes (i.e. recombination “hotspots”; Myers *et al*. 2005), individuals (Kong *et al*., 2004), populations (Dumont *et al*., 2011) and species (Stapley *et al*., 2017).

Genomic studies in humans, cattle, sheep and mice have shown that variation in recombination rate is often heritable, and may have a conserved genetic architecture (Kong *et al*., 2014; Ma *et al*., 2015; Johnston *et al*., 2016; Petit *et al*., 2017). The *loci RNF212, REC8* and *HEI10*, amongst others, have been identified as candidates driving variation in rate, with *PRDM9* driving recombination hotspot positioning in mammals (Baudat *et al*., 2010; Baker *et al*., 2017). This oligogenic architecture suggests that recombination rates and landscapes have the potential to evolve rapidly under different selective scenarios, in turn affecting the rate at which populations respond to selection (Barton and Charlesworth, 1998; Burt, 2000; Otto and Barton, 2001; Gonen *et al*., 2017). However, it remains unclear how representative the above studies are of recombination rate variation and its genetic architecture in natural populations. For example, experimental and domesticated populations tend to be subject to strong selection and have small effective population sizes, both of which have been shown theoretically to indirectly select for increased recombination rates to escape Hill-Robertson interference (Otto and Barton 2001; Otto and Lenormand 2002; but see Munoz-Fuentes *et al*. 2015). Therefore, it may be that prolonged artificial selection results in different recombination dynamics and underlying genetic architectures. As broad recombination patterns are characterised in greater numbers of natural systems (Johnston *et al*., 2016, 2017; Theodosiou *et al*., 2016; Kawakami *et al*., 2017), it is clear that broad and fine-scale recombination rates and landscapes can vary to a large degree even within closely related taxa (Stapley *et al*., 2017). Therefore, determining the genetic architecture of recombination rate in non-model, natural systems are key to elucidating the broad evolutionary drivers of recombination rate variation and quantifying its costs and benefits at the level of the individual.

In this study, we investigate the genetic basis of recombination rate variation in a wild population of Red deer (*Cervus elaphus*) on the island of Rum, Scotland (Clutton-Brock *et al*., 1982). This population has been subject to a long term study since the early 1970s, with extensive pedigree and genotype information for ~2,600 individuals at >38,000 SNPs (Huisman *et al*., 2016; Johnston *et al*., 2017). We use this dataset to identify autosomal crossover rates and their genetic architecture in >1,300 individuals. The aims of the study are to: (a) to determine which common environmental and individual effects, such as age, sex and birth year affect individual recombination rates; (b) to determine if recombination rate is heritable; and (c) to identify genomic regions that are associated with recombination rate variation. Addressing these objectives will provide a foundation for future studies investigating the association between the genetic architecture of recombination rate and individual fitness, to determine how this trait evolves within contemporary natural populations.

## Materials and Methods

### Study population and genomic dataset

The study population of red deer is situated in the North Block of the Isle of Rùm, Scotland (57°02‘N, 6°20‘W) and has been subject to individual monitoring since 1971 (Clutton-Brock *et al*., 1982). Research was conducted following approval of the University of Edinburgh’s Animal Welfare and Ethical Review Body and under appropriate UK Home Office licenses. DNA was extracted from neonatal ear punches, cast antlers and post-mortem tissue (see Huisman *et al*., 2016 for full details). DNA samples from 2880 individuals were genotyped at 50,541 SNP locion the Cervine Illumina BeadChip (Brauning *et al*., 2015) using an Illumina genotyping platform (Illumina Inc., San Diego, CA, USA). SNP genotypes were scored using Illumina GenomeStudio software, and quality control was carried out using the *check.marker* function in GenABEL v1.8-0 (Aulchenko *et al*., 2007) in R v3.3.2, with the following thresholds: SNP genotyping success >0.99, SNP minor allele frequency >0.01, and ID genotyping success >0.99, with 38,541 SNPs and 2,631 IDs were retained. There were 126 pseudoautosomal SNPs identified on the ☓ chromosome (i.e. markers showing autosomal inheritance patterns; Johnston et al., 2017).). Heterozygous genotypes within males at non-pseudoautosomal ☓-linked SNPs were scored as missing. A pedigree of 4,515 individuals has been constructed using microsatellite and SNP data using the software Sequoia (see Huisman, 2017). The genomic inbreeding coefficient 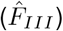, was calculated for each deer in the software GCTA v1.24.3 (Yang *et al*., 2011), using information for all autosomal SNP loci passing quality control. A linkage map of 38,083 SNPs has previously been constructed, with marker orders and estimated base-pair positions known for all 33 autosomes (CEL1 to CEL33) and the ☓ chromosome (CEL34) (Johnston *et al*., 2017 and data archive doi:10.6084/m9.figshare.5002562). All chromosomes are acrocentric with the exception of one metacentric autosome (CEL5).

### Quantification of meiotic crossovers

A standardised sub-pedigree approach was used to identify the positions of meiotic crossovers (Johnston *et al*., 2016). The full pedigree was split as follows: for each focal individual (FID) and offspring pair, a sub-pedigree was constructed that included the FID, its mate, parents and offspring (Figure S1), where all five individuals were genotyped on the SNP chip. This pedigree structure allows phasing of SNPs within the FID, characterising the crossovers occurring in the gamete transferred from the FID to the offspring. All remaining analyses outlined in this section were conducted in the software CRI-MAP v2.504a (Green *et al*., 1990) within the R package crimaptools v0.1 (Johnston *et al*., 2017) implemented in R v3.3.2. Mendelian incompatibilities within sub-pedigrees were identified using the *prepare* function and removed from all affected individuals; sub-pedigrees containing more than 0.1% mismatching locibetween parents and offspring were discarded. The *chrompic* function was used to identify the grand-parental phase of SNP alleles on chromosomes transmitted from the FID to the offspring, and to provide a sex-averaged linkage map. switches in phase indicated the position of a crossover (Figure S1). Individuals with high numbers of crossovers per gamete (>60) were assumed to have widespread phasing errors and were removed from the analysis; the maximum number of crossovers for an individual was 45 in the remaining dataset.

Errors in determining allelic phase can lead to incorrect calling of double crossovers (i.e. ≥ 2 crossovers occurring on the same chromosome) over short map distances. To reduce the likelihood of calling false double crossover events, phased runs consisting of a single SNP were recoded as missing (390 out of 7652 double crossovers; Figure S2) and *chrompic* was rerun. Of the remaining double crossovers, those occurring over distances of ≤ 10cM (as measured by the distance between markers immediately flanking the double crossover) were recoded as missing (170 out of 6959 double crossovers; Figure S2). After this process, 1341 sub-pedigrees were passed quality control, characterising crossovers in gametes transmitted to 482 offspring from 81 unique males and 859 offspring from 256 unique females.

### Genetic architecture of recombination rate variation

#### Heritability and cross-sex genetic correlation

Autosomal crossover count (ACC) was modelled as a trait of the FID. A restricted maximum-likelihood (REML) “animal model” approach (Henderson, 1975) was used to partition phenotypic variance and examine the effect of fixed effects on ACC; these were implemented in ASReml-R (Butler *et al*., 2009) in R v3.3.2. The additive genetic variance was calculated by fitting a genomic relatedness matrix (GRM) constructed for all autosomal markers in GCTA v1.24.3 (Yang *et al*., 2011); the GRM was adjusted assuming similar frequency spectra of genotyped and causal loci using the argument *‐‐grm-adj 0*. There was no pruning of related individuals from the GRM (i.e. we did not use the *‐‐grm-cutoff* argument) as there is substantial relatedness within the population, and initial models included parental effects and common environment which controls for effects of shared environments between relatives. ACC was modelled first using a univariate model:

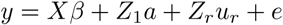

where *y* is a vector of ACC; *X* is an incidence matrix relating individual measures to a vector of fixed effects, *β*; *Z*_1_, and *Z*_*r*_ are incidence matrices relating individual measures with additive genetic and random effects, respectively; *a* and *u*_*r*_ are vectors of GRM additive genetic and additional random effects, respectively; and e is a vector of residual effects. The narrow-sense heritability *h*^2^ was calculated as the ratio of the additive genetic variance to the sum of variance components estimated for all random effects. Model structures were tested with several fixed effects, including sex, 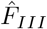 and FID age; random effects included individual identity (i.e. permanent environment) to account for repeated measures in the same FID, maternal and paternal identity, and common environment effects of FID birth year and offspring birth year. The significance of fixed effects was tested with a Wald test, and the significance of random effects was calculated using likelihood-ratio tests (LRT, distributed as *χ*^2^ with 1 degree of freedom) between models with and without the focal random effect. 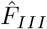 and individual identity were retained in all models, irrespective of statistical significance, to account for possible underestimation of ACC and pseudoreplication, respectively. As the variance in recombination rates differed between the sexes, models were also run for each sex separately. Bivariate models of male and female ACC were run to determine whether additive genetic variation was associated with sex-specific variation and the degree to which this was correlated between the sexes. The additive genetic correlation *r*_A_ was determined using the CORGH error-structure function in ASReml-R (correlation with heterogeneous variances) with *r*_*A*_ set to be unconstrained. Model structure was otherwise the same as for univariate models. To determine whether genetic correlations were significantly different from 0 and 1, the unconstrained model was compared with models where *r*_*A*_ was fixed at values of 0 or 0.999. Differences in additive genetic variance in males and females were tested by constraining both to be equal values using the CORGV error-structure function in ASReml-R. Models then were compared using LRTs with 1 degree of freedom.

#### Genome-wide association study

Genome-wide association studies (GWAS) of ACC were conducted using the function *rGLS* in the R library RepeatABEL v1.1 (Rónnegård *et al*., 2016) implemented in R v3.3.2. This function accounts for population structure by fitting the GRM as a random effect, and allows fitting of repeated phenotypic measures per individual. Models were run including sex and 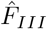 as fixed effects; sex-specific models were also run. Association statistics were corrected for inflation due to population stratification that was not captured by the GRM, by dividing them by the genomic control parameter *λ*, which was calculated as the observed median *χ*^2^ statistic divided by the null expectation median *χ*^2^ statistic (Devlin *et al*., 1999). The significance threshold after multiple testing was calculated using a linkage disequilibrium (LD) based approach in the software *K*_*effective*_ (Moskvina and Schmidt, 2008) specifying a sliding window of 50 SNPs. The effective number of tests was calculated as 35,264, corresponding to a P value of 1.42 × 10^‒06^ at *Ρ* = 0.05. GWAS of ACC included the X chromosome and 458 SNP markers of unknown position. It is possible that some SNPs may show an association with ACC if they are in LD with polymorphic recombination hotspots (i.e. associations in *cis*), rather than SNPs associated with recombination rate globally across the genome (i.e. associations in *trans*). Therefore, we repeated the GWAS modelling *trans* variation only, by examining associations between each SNP and ACC, minus the crossovers that occurred on the same chromosome as the SNP. For example, if the focal SNP occurred on linkage group 1, association was tested with ACC summed over linkage groups 2-33. Marker positions are known relative to the cattle genome vBTA_vUMD_3.1; in cases of significant associations with recombination rate, gene annotations and positions were obtained from Ensembl (Cattle gene build ID BTA_vUMD_3.1.89). LD was calculated between loci in significantly associated regions using the allelic correlation *r*^2^ in the R package LDheatmap v0.99-2 (Shin *et al*., 2006) in R v3.3.2.

#### Regional heritability analysis

As a single locus approach, GWAS has reduced power to detect variants with small effect sizes and/or low linkage disequilibrium with causal mutations (Yang *et al*., 2011). Partitioning additive genetic variance within specific genomic regions (i.e. a regional heritability approach) incorporates haplotype effects and determines the proportion of phenotypic variance explained by defined regions. The additive genetic variance was partitioned across all autosomes in sliding windows of 20 SNPs (with an overlap of 10 SNPs) as follows (Nagamine *et al*., 2012; Bérénos *et al*., 2015):

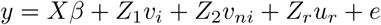

where *y* is a vector of ACC; *X* is an incidence matrix relating individual measures to a vector of fixed effects, *β*; *v* is a vector of additive genetic effects explained by autosomal genomic region in window *i*; *nv* is the vector of the additive genetic effects explained by all autosomal markers not in window *i*; *Z*_1_, and *Z*_2_ are incidence matrices relating individual measures with additive genetic effects for the focal window and the rest of the genome, respectively; *Z*_*r*_ is an incidence matrix relating individual measures with additional random effects, where *u*_*r*_ is a vector of additional random effects; and e is a vector of residual effects. The mean window size was 1.29 ± 0.32 Mb. Models were implemented in ASReml-R (Butler *et al*., 2009) in R v3.3.2. GRMs were constructed in the software GCTA v1.24.3 with the argument *‐‐grm-adj 0* (Yang *et al*., 2011). The significance of additive genetic variance for window i was tested by comparing models with and without the *Z*_1_*v*_*i*_ term with LRT (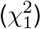). To correct for multiple testing, a Bonferroniapproach was used, taking the number of windows and dividing by 2 to account for window overlap; the threshold P-value was calculated as 2.95 × 10^‒5^ at *α* = 0.05. In the most highly associated region, this analysis was repeated for windows of 20, 10 and 6 SNPs in sliding windows overlapping by *n* – 1 SNPs in order to fine map the associated regions. This was carried out from approximately 5MB before and after the significant region.

#### Accounting for sample size difference between males and females

Sample sizes within this dataset are markedly different between males and females (see above and Table 1). A consequence of this may be that there is lower power to detect associations with male recombination rate. We repeated the heritability and GWAS analyses in sampled datasets of the same size within each sex. Briefly, 482 recombination rate measures (representing the total number in males) were sampled with replacement within the male and female datasets, and the animal model and GWAS analyses were repeated in the sampled dataset. This process was repeated 100 times, with sampling carried out in R v3.3.2. The observed and simulated heritabilities compared to see how often a similar results would be obtained. This was repeated for association at the most highly associated GWAS SNPs and regional heritability regions. The differences between the mean simulated values in each sex were investigated using a Welch two-sample t-test assuming unequal variances.

**Table 1:** Data set information and animal model results for autosomal crossover count (ACC). Numbers in parentheses are the standard error, except for *Mean*, which is the standard deviation. *N*_*OBS*_, *N*_*FID*_ and *N*_*xovers*_ are the number of ACC measures, the number of focal individuals (FIDS) and the total number of crossovers in the dataset. The mean ACC was calculated from the raw data. Vp and VA are the phenotypic variance and additive genetic variance, respectively. *h*^2^, *pe*^2^ and *e*^2^ are the narrow-sense heritability, the permanent environment effect, and the residual effect, respectively; all are calculated as the proportion of *V*_*p*_ that they explain. The additive genetic components were modelled using genomic relatedness matrices. *P*(*h*^2^) is the significance of the *V*_*a*_ term in the model as determined using a likelihood ratio test.

| Analysis | $N_{OBS}$ | $N_{FID}$ | Mean | $N_{xovers}$ | $V_P$ | $V_A$ | $h^2$ | $pe^2$ | $e^2$ | $P(h^2)$ |
| --- | --- | --- | --- | --- | --- | --- | --- | --- | --- | --- |
| Both | 1341 | 337 | 25.03<br>(5.49) | 34911 | 26.42<br>(1.17) | 3.46<br>(1.34) | 0.13<br>(0.05) | 0.05<br>(0.04) | 0.82<br>(0.03) | 0.002 |
| Females | 859 | 256 | 26.62<br>(5.62) | 24025 | 32.02<br>(1.67) | 3.46<br>(1.87) | 0.11<br>(0.06) | 0.05<br>(0.05) | 0.84<br>(0.04) | 0.033 |
| Males | 482 | 81 | 22.21<br>(3.88) | 10886 | 15.33<br>(1.09) | 1.03<br>(1.66) | 0.07<br>(0.11) | 0.06<br>(0.1) | 0.87<br>(0.05) | 0.554 |

#### Haplotyping and effect size estimation

Haplotype construction was carried out to examine haplotype variation within regions significantly associated with recombination rate variation in the regional heritability analysis. SNP data from deer linkage group 12 was phased using SHAPEIT v2.r837 (Delaneau *et al*., 2012), specifying the linkage map positions and recombination rates for each locus. This analysis used pedigree information with the *‐‐duohmm* flag to allow the use of pedigree information in the phasing process (O'Connell *et al*., 2014). Haplotypes were then extracted for the most significant window from the regional heritability analysis.

Effect sizes on ACC for the top GWAS SNPs were estimated using animal models in ASReml-R; SNP genotype was fit as a fixed factor, with pedigree relatedness fit as a random effect to account for the additive genetic variance. To determine the effect sizes on ACC for the regional heritability analysis, animal models were run as follows: for a given haplotype, A, its effect was estimated relative to all other haplotypes combined, i.e. treating them as a single allele, B, by fitting genotypes A/A, A/B and B/B as a fixed factor. This was repeated for each haplotype allele where more than 10 copies were present in the full dataset.

### Data availability

Raw data are publicly archived at doi:10.6084/m9.figshare.5002562 (Johnston *et al*., 2017). Code for the analysis is archived at https://github.com/susjoh/Deer_Recombination_GWAS.

## Results

### Variation and heritability in autosomal crossover count

Autosomal crossover count (ACC) was significantly higher in females than in males, where females had 4.32 ± 0.41 more crossovers per gamete (*Z* = 10.57, *P*_*W ald*_ <0.001; Figure 1); there was no effect of FID age or inbreeding on ACC (P >0.05, Table S1). Females had significantly higher phenotypic variance in ACC than males (*V*_*P*_ = 32.02 and 15.33, respectively; Table 1). ACC was heritable in both sexes combined (*h*^2^ = 0.13, SE = 0.05, P = 0.002) and within females only (*h*^2^ = 0.11, SE = 0.06, P = 0.033), but was not heritable in males (P > 0.05; Table 1). The remaining phenotypic variance was explained by the residual error term, and there was no variance explained by the permanent environment effect, birth year, year of gamete transmission, or parental identities of the FID in any model (animal models P >0.05). Bivariate models of ACC between the sexes indicated that the genetic correlation (*r*_*a*_) between males and females was 0.346, but was not significantly different from zero or one (*P*_*LRT*_ >0.05). This may be due to the relatively small sample size of this dataset resulting in a large standard error around the *r*_*A*_ estimate, or the fact that ACC was not heritable in males. Sampling of 482 measures from each sex showed no difference in heritability estimates between the sexes, indicating reduced power to quantify heritable variation in the smaller male dataset (t = 0.242, P = 0.810, Figure S3).

**Figure 1:**
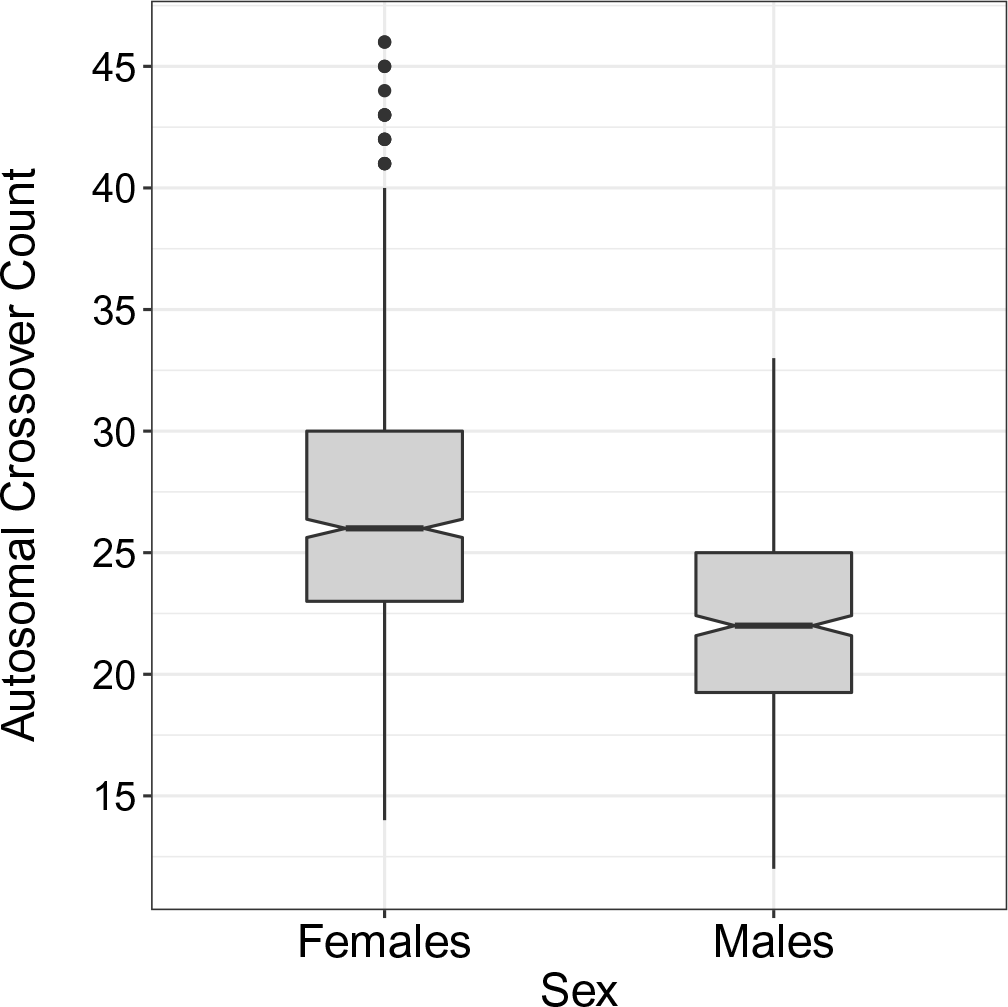
Distribution of ACCs in the raw data for females and males.

### Genetic architecture of autosomal crossover count

#### Genome-wide association study

No SNPs were significantly associated with ACC at the genome-wide level (Figure 2, Tables 2 and S2). The most highly associated SNP in both sexes was *cela1_red_10_26005249* on deer linkage group 12 (CEL12), corresponding to position 26,005,249 on cattle chromosome 10 (BTA10). This marker was also the most highly associated SNP when considering recombination in *trans*, indicating that this region affects ACC across the genome (Table S2). The observed association was primarily driven by female ACC (Table 2, Figure 2). In females, the most highly associated SNP was *cela1_red_10_25661750* on CEL12, corresponding to posiiton 25,661,750 on BTA10. For both SNPs, sampling of 482 measures from each sex showed that the observed associations were significantly higher in females than in males when considering the same sample size (*cela1_red_10_25661750:* t = 18.60, P < 0.001; *cela1_red_10_26005249:* t = 4.89, P < 0.001; Figure S4). Based on its position relative to the cattle genome, *cela1_red_10_26005249* was ~600bp upstream of an olfactory receptor *OR5AU1* and ~24kb downstream from a gene of unknown function (ENSB-TAG00000011396). There were four candidate genes within 1Mb of both loci, including *TOX4*, *CHD8*, *SUPT16H* and *CCNB1IP1* (Figure 4; see Discussion). Similar results were obtained when considering recombination rate on all chromosomes excluding that on which the SNP occurred, indicating that all associations affect recombination rate variation in *trans* across the genome (Table S2).

**Figure 2:**
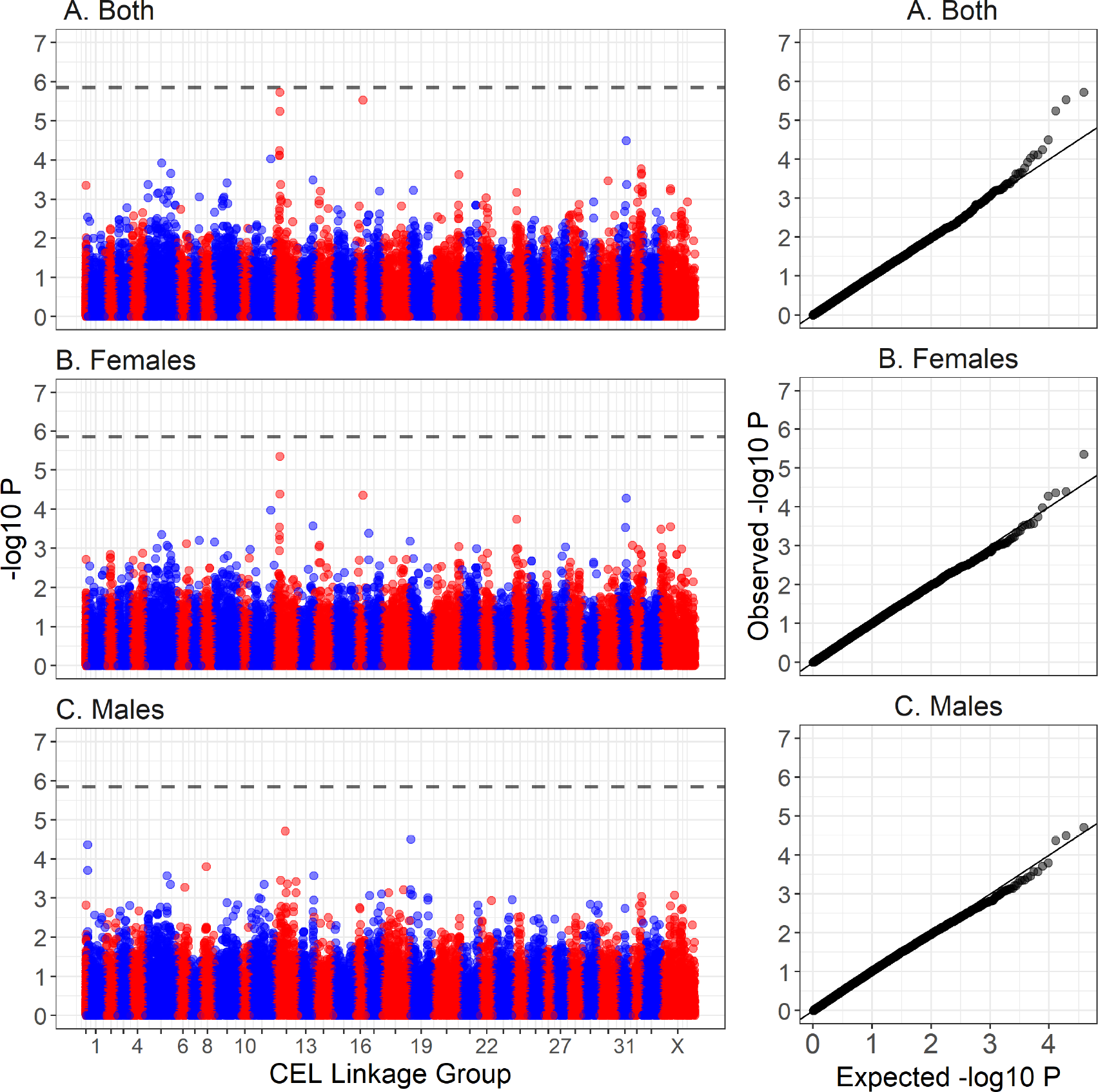
Manhattan plot of genome-wide association of autosomal crossover count (ACC) for (A) all deer, (B) females only and (C) males only. The dashed line is the genome-wide significance threshold equivalent to P <0.05. The left-hand plots show association relative to the estimated genomic positions on deer linkage groups from Johnston *et al*. (2017). Points have been colour coded by chromosome. The right-hand plots show the distribution of observed *‒log*_10_*P* values against those under the null expectation. Association statistics have been corrected for the genomic control inflation factor *λ*. Underlying data are provided in Table S2 and sample sizes are given in Table 1.

**Table 2:** The top five most significant hits from a genome-wide association study of ACC in (A) Both sexes, (B) Females only and (C) Males only. No SNPs reached the genome-wide significance of P = 1.42 × 10^‒06^. The SNP locus names indicate the position of the SNPs relative to the cattle genome assembly vBTA_vUMD_3.1 (indicated by *Chromosome_Position*). Linkage groups and map positions (in centiMorgans, cM) are from Johnston *et al*. (2017). A and B are the reference alleles. Effect B is the estimated effect and standard error of the B allele as estimated in RepeatABEL (RönnegÅrd *et al*., 2016). P-values have been corrected for the genomic inflation parameter *λ*. Full results are available in Table S2.

| Sex | SNP Locus | Deer Linkage Group | Map Position (cM) | A | B | Effect B | (SE) | $\chi^2_1$ | *P* | MAF |
| --- | --- | --- | --- | --- | --- | --- | --- | --- | --- | --- |
| A. Both | cela1_red_10_26005249 | 12 | 36.4 | G | A | 1.53 | 0.28 | 22.73 | 1.87e-06 | 0.33 |
|  | cela1_red_8_100681301 | 16 | 43.5 | A | G | 6.42 | 1.19 | 21.87 | 2.91e-06 | 0.02 |
|  | cela1_red_10_25661750 | 12 | 35.6 | A | G | 2.18 | 0.42 | 20.6 | 5.67e-06 | 0.1 |
|  | cela1_red_1_35423049 | 31 | 46.2 | A | G | 1.4 | 0.29 | 17.31 | 3.18e-05 | 0.25 |
|  | cela1_red_10_21372438 | 12 | 34.5 | A | G | 1.22 | 0.26 | 16.2 | 5.69e-05 | 0.42 |
| B. Females | cela1_red_10_25661750 | 12 | 35.6 | A | G | 2.81 | 0.56 | 21.07 | 4.44e-06 | 0.1 |
|  | cela1_red_10_26005249 | 12 | 36.4 | G | A | 1.58 | 0.35 | 16.84 | 4.07e-05 | 0.33 |
|  | cela1_red_8_100681301 | 16 | 43.5 | A | G | 6.25 | 1.4 | 16.72 | 4.34e-05 | 0.02 |
|  | cela1_red_1_35423049 | 31 | 46.2 | A | G | 1.62 | 0.37 | 16.37 | 5.22e-05 | 0.25 |
|  | cela1_red_11_91378678 | 11 | 86.5 | A | G | 13.61 | 3.22 | 15.03 | 1.06e-04 | 0.02 |
| C. Males | cela1_red_10_49732924 | 12 | 52.6 | G | A | -2.9 | 0.66 | 18.25 | 1.94e-05 | 0.14 |
|  | cela1_red_1_128593904 | 19 | 13.5 | G | A | -1.99 | 0.46 | 17.32 | 3.15e-05 | 0.18 |
|  | cela1_red_15_6941417 | 1 | 8.9 | A | G | -1.93 | 0.46 | 16.74 | 4.28e-05 | 0.21 |
|  | cela1_red_2_101879999 | 8 | 35.2 | G | A | 1.51 | 0.39 | 14.28 | 1.58e-04 | 0.44 |
|  | cela1_red_15_7417500 | 1 | 9.2 | A | C | -1.93 | 0.5 | 13.9 | 1.93e-04 | 0.17 |

#### Regional heritability analysis

The genome-wide regional heritability analysis of ACC showed a significant association in both sexes and in females only with a ‐;2.94Mb region on CEL12 (Figure 3, Table 3). The most highly associated window (~1.36 Mb) within this region contained 42 genes, including REC8 meiotic recombination protein (*REC8;* 20,810,610 - 20,817,662 bp on BTA10). Detailed examination of this region in sliding windows of 6, 10 and 20 SNPs found the highest association at a 10 SNP window of ~463kb containing 36 genes, including *REC8* (Table 3). This region explained all heritable variation in recombination rate, with regional heritability estimates of 0.143 (SE = 0.053) and 0.146 (SE = 0.045) for all deer and females only, respectively. The sex-specific effect was supported by sampling of 482 measures, where females had consistently higher associations than in males (t = 19.03, P < 0.001, Figure S5). The total significant region after detailed examination was ~3.01Mb wide, flanked by SNPs *cela1_red_10_18871213* and *cela1_red_10_21878407* (Figure 4 & Table S4) and containing ~87 genes. This wider region contained the protein coding region for ring finger protein 212B (*RNF212B;* 21,466,337 - 21,494,696 bp on BTA10), a homologue of *RNF212*, which has been directly implicated in synapsis and crossing-over during meiosis in mice (Reynolds *et al*., 2013). Genetic variants at both *RNF212B* and *RNF212* have been associated with recombination rate variation in humans, cattle and sheep (Kong *et al*., 2008; Ma *et al*., 2015; Johnston *et al*., 2016; Petit *et al*., 2017). Whilst this region was close to the most highly associated SNPs from the genome-wide association study, there was no overlap between the two analyses, with the mostly highly associated regions separated by an estimated ~5.5Mb (Figure 4). The mean *r*^2^ LD between the top regional heritability window and the top GWAS SNPs was 0.258 for *cela1_red_10_25661750* and 0.276 for *cela1_red_10_26005249*, with the top *r*^2^ of 0.665 observed between the SNPs *cela1_red_10_21807996* and *cela1_red_10_26005249* (Figure 4).

**Figure 3:**
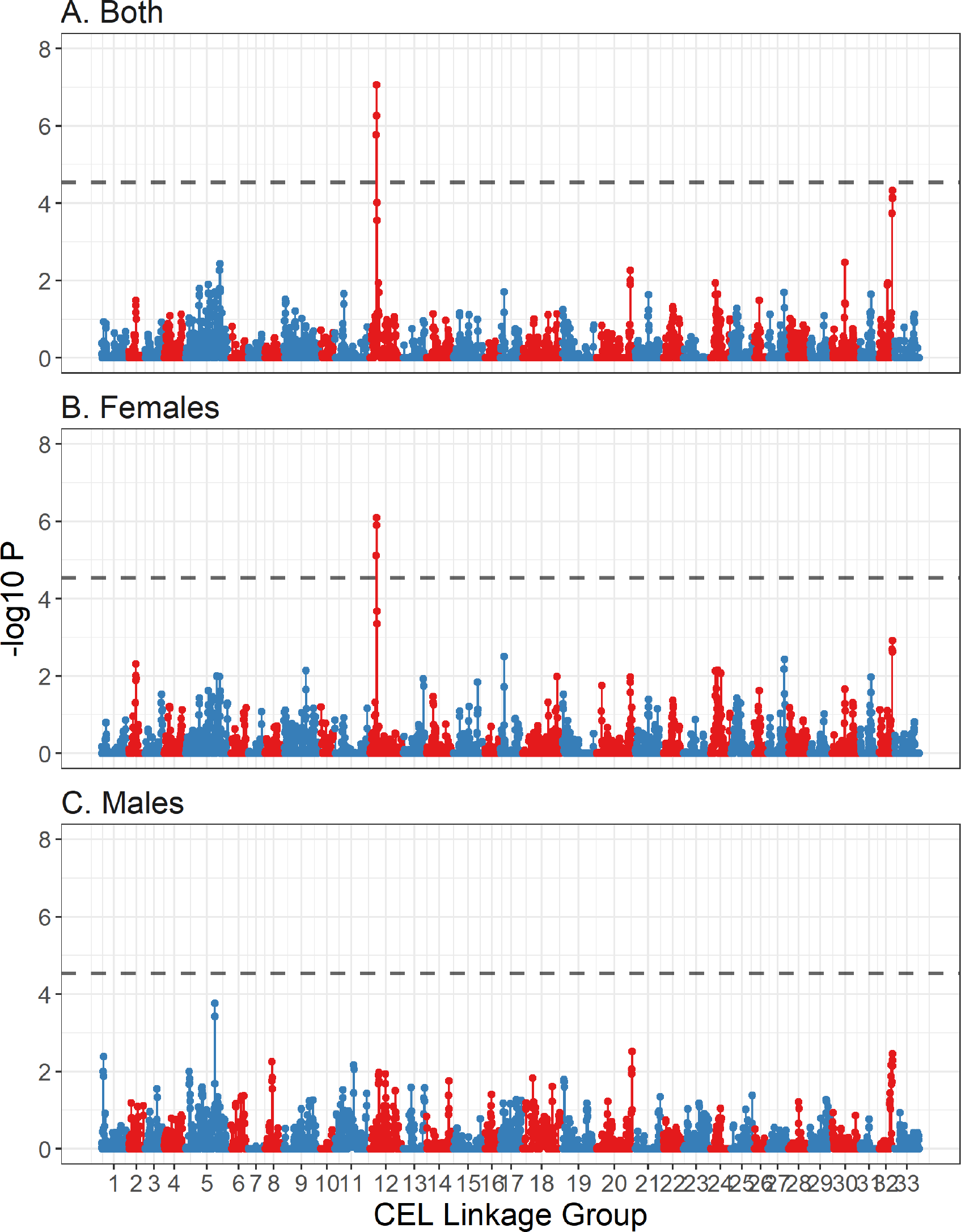
Regional heritability plot of association of autosomal crossover count for (A) all deer, (B) females only and (C) males only. Each point represents a sliding window of 20 SNPs with an overlap of 10 SNPs. The dashed line is the genome-wide significance threshold equivalent to P <0.05 as calculated using Bonferroni. Lines have been colour coded by chromosome. Underlying data are provided in Table 3.

**Figure 4:**
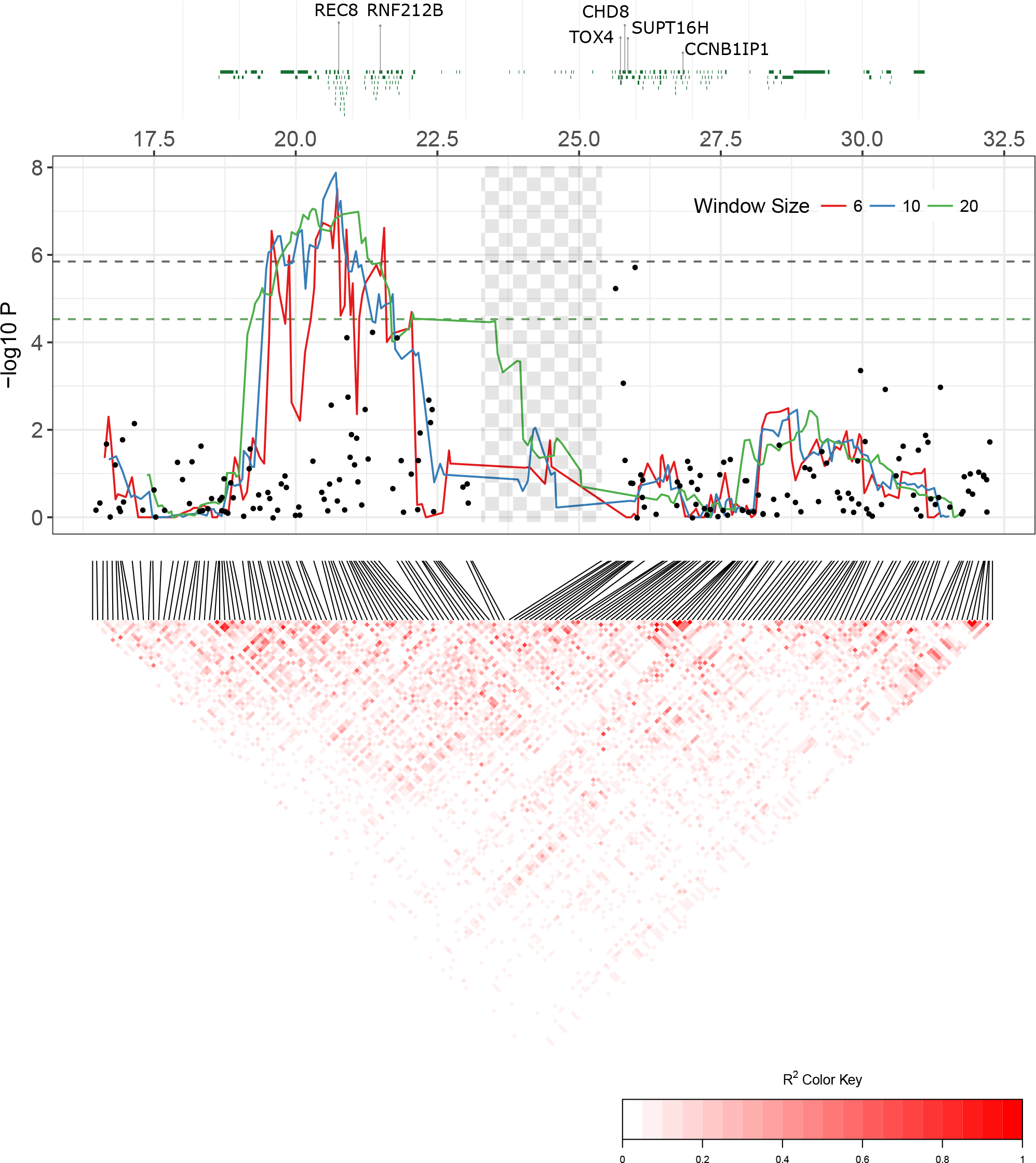
Detailed figure of genes, association statistics and linkage disequilibrium patterns at the most highly associated region on on CEL12 (homologous to BTA10) for all deer of both sexes. All X-axis positions are given relative to the cattle genome vBTA_vUMD_3.1. The top panel shows protein coding regions, with annotation for candidate loci. The central panel shows the results for the regional heritability analysis (where lines represents a sliding windows of 6,10 and 20 SNPs with an overlap of n-1 SNPs) and the genome-wide association study (where points indicate single SNP associations). The dashed lines are the genome-wide significance thresholds (green = regional heritability, black = genome-wide association). The checked shaded area shows the position of the T cell receptor alpha/delta locus (see Discussion). Underlying data are provided in Table S4. The lower panel shows linkage disequilibrium between each loci using allelic correlations (*r*^2^).

**Table 3:** The most significant hits from a regional heritability analysis of ACC in (A) Both sexes, (B) Females only and (C) Males only. Sliding windows were 20 SNPs wide with an overlap of 10 SNPs. Lines in italics are the most highly associated regions from detailed examination of significant regions - in each case these are for 10 SNP windows. The *χ*^2^ and P values are for likelihood ratio test comparisons between models with and without a genomic relatedness matrix for that window; values in bold type are significant the the genome-wide level. The SNP locus names indicate the position of the SNPs relative to the cattle genome assembly vBTA_vUMD_3.1 (indicated by *Chromosome_Position*). Full results are available in Tables S3 & S4.

| Sex | Deer Linkage Group | $\chi^2_1$ | P | First SNP | Last SNP | Region $h^2$ | SE |
| --- | --- | --- | --- | --- | --- | --- | --- |
| A. Both | *12* | *32.30* | *1.32e-08* | *cela1_red_10_20476277* | *cela1_red_10_20939342* | *0.143* | *0.053* |
|  | 12 | 28.62 | 8.81e-08 | cela1_red_10_19617695 | cela1_red_10_20977030 | 0.080 | 0.043 |
|  | 12 | 25.11 | 5.41e-07 | cela1_red_10_20519507 | cela1_red_10_21807996 | 0.080 | 0.045 |
|  | 12 | 22.91 | 1.70e-06 | cela1_red_10_18871213 | cela1_red_10_20476277 | 0.105 | 0.055 |
|  | 32 | 16.55 | 4.73e-05 | cela1_red_27_38731584 | cela1_red_27_40264086 | 0.056 | 0.034 |
|  | 32 | 15.76 | 7.21e-05 | cela1_red_27_39821973 | cela1_red_27_41274975 | 0.071 | 0.045 |
| B. Females | *12* | *28.14* | *1.13e-07* | *cela1_red_10_20476277* | *cela1_red_10_20939342* | *0.146* | *0.045* |
|  | 12 | 24.34 | 8.06e-07 | cela1_red_10_19617695 | cela1_red_10_20977030 | 0.089 | 0.048 |
|  | 12 | 23.5 | 1.25e-06 | cela1_red_10_20519507 | cela1_red_10_21807996 | 0.102 | 0.056 |
|  | 12 | 20.03 | 7.61e-06 | cela1_red_10_18871213 | cela1_red_10_20476277 | 0.133 | 0.068 |
|  | 12 | 13.72 | 2.12e-04 | cela1_red_10_21000545 | cela1_red_10_22450693 | 0.089 | 0.054 |
|  | 12 | 12.32 | 4.49e-04 | cela1_red_10_21878407 | cela1_red_10_26041475 | 0.177 | 0.087 |
| C. Males | 5 | 14.07 | 1.76e-04 | cela1_red_19_15289588 | cela1_red_19_16108226 | 0.133 | 0.052 |
|  | 5 | 12.61 | 3.84e-04 | cela1_red_19_15753501 | cela1_red_19_16923111 | 0.137 | 0.058 |
|  | 20 | 8.77 | 3.06e-03 | cela1_red_3_110763634 | cela1_red_3_112123206 | 0.142 | 0.085 |
|  | 32 | 8.51 | 3.52e-03 | cela1_red_27_38731584 | cela1_red_27_40264086 | 0.119 | 0.076 |
|  | 1 | 8.21 | 4.17e-03 | cela1_red_15_6354196 | cela1_red_15_7482634 | 0.123 | 0.056 |

A second region on linkage group 32 almost reached genome-wide significance in the regional heritability analysis, corresponding to the region ‐38.7 - 41.3Mb on cattle chromosome 27. This region contained the locus topoisomerase (DNA) II beta (*TOP2B*); inhibitors of this gene lead to defects chromosome segregation and heterochromatin condensation during meiosis I in mice, *Drosophila melanogaster* and *Caenorhabditis elegans* (Li *et al*. 2013; Gómez *et al*. 2014; Hughes and Hawley 2014; Jaramillo-Lambert *et al*. 2016; Figure 3, Tables 3 and S3). Full results for the regional heritability analyses are provided in Tables S3 and S4.

#### Effect size estimation

At most highly associated GWAS SNP, *cela1_red_10_26005249*, carrying one or two copies of the G allele conferred 3.3 to 3.9 fewer crossovers per gamete in females (Wald P < 0.001) and 1.8 - 2.8 fewer crossovers per gamete in males (P = 0.009; Table 4). The most highly associated SNP in females, *cela1_red_10_25661750*, had a significant effect on ACC in females (P < 0.001) but not in males (P > 0.05; Table 4). This locus conferred 2.03 more crossovers in A/G females and 13.68 more in G/G females; however, the latter category contained 7 unique measures in only two individuals, and so this estimate is likely to be subject to large sampling error.

**Table 4:** Effect sizes for the most highly associated GWAS SNPs and for the AGGAGAGAAG haplotype at the most highly associated regional heritability region. Models were run for each sex separately and included a pedigree relatedness as a random effect. Count and ID Count indicate the number of ACC measures and the number of unique individuals for each genotype, respectively. Wald.P indicates the P-value for a Wald test of genotype as a fixed effect.

| Locus | Sex | Genotype | Count | ID Count | Solution | S.E. | Z Ratio | Wald.P |
| --- | --- | --- | --- | --- | --- | --- | --- | --- |
| cela1_red_10_26005249 | Female | A/A (Intercept) | 98 | 28 | 29.575 | 0.732 | 40.43 | 3.43e-06 |
|  |  | A/G | 388 | 114 | -3.269 | 0.733 | -4.46 |  |
|  |  | G/G | 377 | 116 | -3.888 | 0.786 | -4.944 |  |
|  | Male | A/A (Intercept) | 27 | 6 | 24.405 | 0.964 | 25.327 | 8.90e-03 |
|  |  | A/G | 248 | 40 | -1.813 | 0.993 | -1.826 |  |
|  |  | G/G | 207 | 35 | -2.863 | 1.025 | -2.793 |  |
| cela1_red_10_25661750 | Female | A/A (Intercept) | 688 | 208 | 25.979 | 0.36 | 72.114 | 6.34e-10 |
|  |  | A/G | 168 | 48 | 2.026 | 0.56 | 3.619 |  |
|  |  | G/G | 7 | 2 | 13.684 | 2.386 | 5.736 |  |
|  | Male | A/A (Intercept) | 411 | 65 | 22.13 | 0.367 | 60.345 | 0.399 |
|  |  | A/G | 57 | 14 | 0.934 | 0.69 | 1.353 |  |
|  |  | G/G | 14 | 2 | 0.345 | 1.512 | 0.228 |  |
| Haplotype<br>AGGAGAGAAG | Female | A/A (Intercept) | 690 | 208 | 25.905 | 0.364 | 71.125 | 7.29e-09 |
|  |  | A/B | 160 | 46 | 2.387 | 0.573 | 4.166 |  |
|  |  | B/B | 13 | 4 | 8.701 | 1.73 | 5.029 |  |
|  | Male | A/A (Intercept) | 406 | 66 | 22.017 | 0.36 | 61.153 | 0.037 |
|  |  | A/B | 62 | 13 | 1.72 | 0.669 | 2.571 |  |
|  |  | B/B | 14 | 2 | 0.506 | 1.492 | 0.339 |  |
| Haplotype<br>AGAGAAGAGA | Female | A/A (Intercept) | 795 | 242 | 26.591 | 0.451 | 58.928 | 0.026 |
|  |  | A/B | 68 | 16 | -2.244 | 1.005 | -2.233 |  |
|  | Male | A/A (Intercept) | 481 | 80 | 22.279 | 0.351 | 63.403 | 0.775 |
|  |  | A/B | 1 | 1 | -1.122 | 3.925 | -0.286 |  |

Atotal of 17 haplotypes in the 10 SNP region spanning *cela1_red_10_20476277* and *cela1_red_1* had more than ten copies in unique individual females (Table S5). Of these, two haplotypes, AGGAGAGAAG and AGAGAAGAGA, had a significant effect on ACC relative to all other haplotypes (Tables 4 and S5, Figure S6). Haplotype AGGAGAGAAG increased female ACC by 2.4 crossovers per gamete in heterozygotes (P < 0.001); homozygotes for the haplotype were rare (13 measures in 4 individuals) and so the large effect size estimate was again likely to be subject to large sampling effects (Table 4). The haplotype AGAGAAGAGA reduced female ACC by 2.2 crossovers per gamete in heterozygous individuals (P < 0.05; Table 4). The *r*^2^ LD between haplotype AGGAGAGAAG and the two most highly associated GWAS SNPs was 0.464 and 0.885 for *cela1_red_10_26005249* and *cela1_red_10_25661750*, respectively; for haplotype AGAGAAGAGA, it was 0.229 and 0.036 for *cela1_red_10_26005249* and *cela1_red_10_25661750*, respectively.

## Discussion

In this study, we have shown that autosomal crossover count (ACC) is 1.2× higher in females than in males, with females exhibiting higher phenotypic and additive genetic variance for this trait; ACC was not significantly heritable in males. Almost all genetic variation in females was explained by a ~7Mb region on deer linkage group 12. This region contained several candidate genes, including *RNF212B* and *REC8*, which have previously been implicated in recombination rate variation in other mammal species, including humans, mice, cattle and sheep (Kong *et al*., 2008; Reynolds *et al*., 2013; Ma *et al*., 2015; Johnston *et al*., 2016; Petit *et al*., 2017). Here, we discuss in detail the genetic architecture of individual recombination rate, candidate genes underlying heritable variation, sexual-dimorphism in this trait and its architecture, and the conclusions and implications of our findings for other studies of recombination in the wild.

### The genetic architecture of individual recombination rate

Using complementary trait mapping approaches, we identified a ~7Mb region on deer linkage group 12 (homologous to cattle chromosome 10) associated with ACC. The most highly associated GWAS region occurred at ‐25.6 and 26Mb (relative to the cattle genome position), although this association was not significant at the genome-wide level. The most highly associated regional heritability region occurred between ‐20.5 and 20.9MB, around 5Mb away from the top GWAS hits (Figure 4); association at this region was significant at the genome-wide level and explained almost all of the heritable variation in ACC in both sexes and in females only. Most variation in mean ACC was attributed to two haplotypes within this region (Tables 4 and S5; Figure S6).

At present, it is not clear why the results of the two analyses occur in close vicinity, yet do not overlap. Assuming homology with humans, cattle, sheep and mice (Ensembl release 91, Zerbino *et al*. 2018), the two regions are separated by the highly repetitive T-cell receptor alpha/delta variable (TRAV/DV) locus, which may contain up to 400 TRAV/DV genes in cattle (Reinink and Van Rhijn 2009; Figure 4). This region is of an unknown size in deer; relative to the cattle genome, these regions are separated by 4.72Mb, but the deer linkage map distance is estimated as 1.86 centiMorgans (cM). The sex-averaged genome-wide recombination rate in deer is ‐1.04cM/Mb, suggesting this genomic region may be shorter in deer (Johnston *et al*., 2017) and that these two regions are in closer vicnity. This is supported by both the linkage map distance and patterns of linkage disequilibrium between the associated loci, particularly at the associated haplotypes (see Results & Figure 4). In addition, the small sample size used in the current study may result in increased sensitivity to sampling effects and bias in the estimation of the relative contribution of SNPs to the trait mean (GWAS) or variance (Regional heritability). Further investigation with higher samples sizes, whole genome sequencing approaches and improved genome assembly may allow more accurate determination of the most likely candidate genes and potential causal mutations (coding or regulatory) within this species.

### Candidate genes for recombination rate variation

#### Regional heritability analysis

The most highly associated region in the regional heritability analysis contained the gene *REC8*, the protein of which is required for the separation of sister chromatids during meiosis (Parisi *et al*., 1999). It also contained *RNF212B*, a paralogue of *RNF212*; the latter has been associated with recombination rate variation in humans, cattle and sheep (Kong *et al*., 2008; Sandor *et al*., 2012; Johnston *et al*., 2016; Petit *et al*., 2017), with the *REC8/RNF212B* region is associated with recombination rate in cattle, and has a large effect size on ACC phenotype than RNF212 in this species (Sandor *et al*., 2012; Ma *et al*., 2015). A second region on deer linkage group 32 almost reached genome-wide significance (Figure 3, Tables 3 and S3). This region was relatively genepoor, but contained ‐6 genes, including the candidate topoisomerase (DNA) II beta (*TOP2B*): inhibitors of this gene lead to defects in chromosome segregation and heterochromatin condensation during meiosis I in mice, *Drosophila melanogaster* and *Caenorhabditis elegans* (Li *et al*., 2013; Gómez *et al*., 2014; Hughes and Hawley, 2014; Jaramillo-Lambert *et al*., 2016). No association was observed at the region homologous to *RNF212* (predicted to be at position ~109.2Mb on cattle chromosome 6, corresponding to ~57.576cM on deer linkage group 6) for the GWAS or regional heritability analysis.

#### Genome-wide association study (GWAS)

Examination of annotated regions within 500kb of either side of the most significant GWAS SNPs identified three genes, TOX High Mobility Group Box Family Member 4 (*TOX4*), Chromodomain Helicase DNA Binding Protein 8 (*CHD8*) and SPT16 Homologue Facilitates Chromatin Remodelling Subunit (*SUPT16H*). These genes are involved in with chromatin binding and structure (*SP16H*, *TOX4*), histone binding (*CHD8*, *SUPT16H*), nucleosome organisation (*SP16H*) and cell cycle transition (TOX4). One of these genes, *SUPT16H*, interacts with NIMA related kinase 9 (*NEK9*), which is involved with meiotic spindle organisation, chromosome alignment and cell cycle progression in mice (Yang *et al*., 2012) and is a strong candidate locus for crossover interference in cattle (Wang *et al*., 2016). The SNP *cela1_red_10_26005249* was —825kb from Cyclin B1 Interacting Protein 1 (*CCNB1IP1*), also known as Human Enhancer Of Invasion 10 (*HEI10*), which interacts with *RNF212* to allow recombination to progress into crossing-over in mice (Qiao *et al*., 2014) and *Arabidopsis* (Chelysheva *et al*., 2012); this locus is also associated with recombination rate variation in humans (Kong *et al*., 2014).

#### Sexual dimorphism in genetic architecture of recombination rate

The results of this analysis suggest that there is sexual dimorphism in the genetic architecture of recombination rate variation in deer. Higher female recombination rates are typical in mammals (Brandvain and COOP, 2012) and in this system is driven by higher female recombination rates near centromeric regions (Johnston et al., 2017). Male ACC was not significantly heritable, although we could not rule out that this was a consequence of the smaller sample size relative to females (Figure S3). No regions of the genome were significantly associated with male ACC in the regional heritability and GWAS analyses, but sampling did indicate that differences observed between male and female genomic associations were (Figures S4 & S5). Investigation of genetic correlations between males and females was inconclusive, as the *r*_*A*_ of ACC was not significantly different from 0 or 1. The observed sex differences are consistent with previous studies of the genetic architecture of ACC in mammals, where a sexually-dimorphic architecture has been observed at the paralogous *RNF212* region in humans and sheep (Kong *et al*., 2014; Johnston *et al*., 2016). Nevertheless, some observed associations were stronger when considering both male and female deer in the same analysis, for example at the most highly associated GWAS SNP, and the amplified signal for the regional heritability analysis on linkage group 33 (Figures 2 & 3), suggesting that there may be some degree of shared architecture within these regions.

#### Conclusions and implications for studies of recombination in the wild

We have shown that recombination rate is heritable in female red deer, and that it has a sexually dimorphic genetic architecture. Genomic regions associated with recombination rate in red deer are as sociated with this trait in other mammal species, supporting the idea that recombination rate variation has a conserved genetic architecture across distantly related taxa. A key motivation for this study is to compare how recombination rate and its genetic architecture is similar or different to that of model species that have experienced strong selection in their recent history, such as humans, cattle, mice and sheep. The heritability of recombination rate in deer was lower than that observed in other mammal systems (Dumont *et al*., 2009; Kong *et al*., 2014; Ma *et al*., 2015; Johnston *et al*., 2016), with no observed heritable variation present in male deer. Whilst we were able to test their effects, we found no contribution of contribution of individual and common environmental effects on recombination rate (i.e. age, year of birth, year of gamete transmission); indeed, most phenotypic variance in recombination was attributed to residual effects. This suggests that despite some underlying genetic variation, recombination rate is mostly driven by stochastic effects, or otherwise unmeasured effects.

This represents one of the smallest datasets in which recombination rate has been investigated, and so it may be that the observed effects are underestimated due to the small sample size, sampling effects, or perhaps that other genetic variants present in this species do not segregate in the Rùm deer population. Nevertheless, identification of clear candidate genes and their effects on phenotype represents a valuable contribution to understanding the genetic of recombination more broadly. Ultimately, our findings allow future investigation of the fitness consequences of variation in recombination rate and the relationship between identified variants and individual life-history variation, to address questions on the maintenance of genetic variation for recombination rates, and the relative roles of selection, sexually antagonistic effects and stochastic processes in contemporary natural populations.

## Acknowledgements

We thank T. Clutton-Brock and L. Kruuk for maintaining the long-term field project, and A. Morris, S. Morris, M. Baker, F. Guinness and many others for collecting field data and DNA samples. We thank Scottish Natural Heritage for permission to work on the Isle of Rùm National Nature Reserve. We thank P. Ellis for sample preparation and DNA extraction, and the Wellcome Trust Clinical Research Facility Genetics Core in Edinburgh for performing the genotyping. This work has made extensive use of the resources provided by the University of Edinburgh Compute and Data Facility (http://www.ecdf.ed.ac.uk/). The long-term project on Rùm red deer is funded by the UK Natural Environment Research Council, and SNP genotyping was supported by a European Research Council Advanced Grant to J.M.P‥ S.E.J. is supported by a Royal Society University Research Fellowship.

## Author Contributions

S.E.J and J.M.P. conceived the study. J.M.P organised the collection of samples. J.H. conducted DNA sample extraction and genotyping and constructed the pedigree. S.E.J. analysed the data and wrote the paper. All authors contributed to revisions.

## References

1. Arbeithuber, B., A. J. Betancourt, T. Ebner, and I. Tiemann-Boege, 2015 Crossovers are associated with mutation and biased gene conversion at recombination hotspots. Proc. Natl. Acad. Sci. U.S.A. 112: 2109-2114.

2. Aulchenko, Y. S., S. Ripke, A. Isaacs, and C. M. van Duijn, 2007 GenABEL: an R library for genome-wide association analysis. Bioinformatics 23: 1294-1296.

3. Baker, Z., M. Schumer, Y. Haba, L. Bashkirova, C. Holland, etal., 2017 Repeated losses of PRDM9-directed recombination despite the conservation of PRDM9 across vertebrates. eLife 6: e24133.

4. Barton, N. H., and B. Charlesworth, 1998 Why sex and recombination? Science 74: 187-195.

5. Battagin, M., G. Gorjanc, A.-M. Faux, S. E. Johnston, and J. M. Hickey, 2016 Effect of manipulating recombination rates on response to selection in livestock breeding programs. Genet. Select. Evol. 48: 44.

6. Baudat, F., J. Buard, C. Grey, A. Fledel-Alon, C. Ober, etal., 2010 PRDM9 is a major determinant of meiotic recombination hotspots in humans and mice. Science 327: 836-840.

7. Bérénos, C., P. A. Ellis, J. G. Pilkington, S. H. Lee, J. Gratten, etal., 2015 Heterogeneity of genetic architecture of body size traits in a free-living population. Mol. Ecol. 24: 1810-1830.

8. Brauning, R., P. J. Fisher, A. F. McCulloch, R. J. Smithies, J. F. Ward, etal., 2015 Utilization of high throughput genome sequencing technology for large scale single nucleotide polymorphism discovery in red deer and canadian elk. bioRxiv doi:10.1101/027318.

9. Burt, A., 2000 Sex, Recombination, and the Efficacy of Selection - was Weismann Right? Evolution 54: 337-351.

10. Butler, D. G., b. R. Cullis, A. R. Gilmour, and B. J. Gogel, 2009 Mixed Models for S language Environments: ASReml-R reference manual.

11. Chelysheva, L., D. Vezon, A. Chambon, G. Gendrot, L. Pereira, etal., 2012 The Ara-bidopsis HEI10 Is a New ZMM Protein Related to Zip3. PLoS Genet. 8: e1002799.

12. Clutton-Brock, t., F. Guinness, and S. Albon, 1982 Red Deer. Behaviour and Ecology of Two Sexes‥ University of Chicago Press.

13. Delaneau, O., J.-F. Zagury, and J. Marchini, 2012 Improved whole-chromosome phasing for disease and population genetic studies. Nat. Methods 10: 5-6.

14. Devlin, A. B., K. Roeder, and B. Devlin, 1999 Genomic Control for Association. Biometrics 55:997-1004.

15. Dumont, b. L., K. W. Broman, and B. A. Payseur, 2009 Variation in genomic recombination rates among heterogeneous stock mice. Genetics 182: 1345-9.

16. Dumont, b. L., M. A. White, b. Steffy, T. Wiltshire, and B. A. Payseur, 2011 Extensive recombination rate variation in the house mouse species complex inferred from genetic linkage maps. Genome Res. 21: 114-125.

17. Felsenstein, J., 1974 The evolutionary advantage of recombination. Genetics 78: 737-756.

18. Gómez, R., A. Viera, I. Berenguer, E. Llano, A. M. Pendâs, et al., 2014 Cohesin removal precedes topoisomerase IIa-dependent decatenation at centromeres in male mammalian meiosis II. Chromosoma 123: 129-146.

19. Gonen, S., M. Battagin, S. Johnston, G. Gorjanc, and J. Hickey, 2017 The potential of shifting recombination hotspots to increase genetic gain in livestock breeding. Genet. Select. Evol. 49: 55.

20. Green, P., K. Falls, and S. Crooks, 1990 Documentation for CRIMAP, version 2.4. Washington university School of Medicine.

21. Hassold, T., and P. Hunt, 2001 To err (meiotically) is human: the genesis of human aneuploidy. Nat. Rev. Genet. 2: 280-291.

22. Henderson, C. R., 1975 Best linear unbiased estimation and prediction under a selection model. Biometrics 31: 423-447.

23. Hill, W. G., and A. Robertson, 1966 The effect of linkage on limits to artificial selection. Genet. Res. 8: 269-294.

24. Hughes, S. E., and R. S. Hawley, 2014 Topoisomerase II Is Required for the Proper Separation of Heterochromatic Regions during Drosophila melanogaster Female Meiosis. PLoS Genet. 10: e1004650.

25. Huisman, J., 2017 Pedigree reconstruction from SNP data: parentage assignment, sibship clustering and beyond. Mol. Ecol. Resour. 17: 1009-1024.

26. Huisman, J., L. E. B. Kruuk, P. A. Ellis, T. Clutton-Brock, and J. M. Pemberton, 2016 Inbreeding depression across the lifespan in a wild mammal population. Proc. Natl. Acad. Sci. U.S.A. 113: 3585-3590.

27. Inoue, K., and J. R. Lupski, 2002 Molecular mechanisms for genomic disorders. Ann. Rev. Genomics Hum. Genet. 3: 199-242.

28. Jaramillo-Lambert, A.., A. S. Fabritius, T. J. Hansen, H. E. Smith, and A. Golden, 2016 The Identification of a Novel Mutant Allele of topoisomerase II in Caenorhabditis elegans Reveals a Unique Role in Chromosome Segregation During Spermatogenesis. Genetics 204:1407-1422.

29. Johnston, S. E., C. Bérénos, J. Slate, and J. M. Pemberton, 2016 Conserved genetic architecture underlying individual recombination rate variation in a wild population of soay sheep (ovis aries). Genetics 203: 583-598.

30. Johnston, S. E., J. Huisman, P. A. Ellis, and J. M. Pemberton, 2017 A High Density Linkage Map Reveals Sexual Dimorphism in Recombination Landscapes in Red Deer (*Cervus elaphus*). G3 (Bethesda) 7: 2859-2870.

31. Kawakami, t., C. F. Mugal, A. Suh, A. Nater, R. Burri, etal., 2017 Whole-genome patterns of linkage disequilibrium across flycatcher populations clarify the causes and consequences of fine-scale recombination rate variation in birds. Mol. Ecol. 26: 4158-4172.

32. Kong, A., J. Barnard, D. F. Gudbjartsson, G. Thorleifsson, G. Jonsdottir, etal., 2004 Recombination rate and reproductive success in humans. Nat. Genet. 36: 1203-1206.

33. Kong, A., G. Thorleifsson, M. L. Frigge, G. Masson, D. F. Gudbjartsson, etal., 2014 Common and low-frequency variants associated with genome-wide recombination rate. Nat. Genet. 46: 11-6.

34. Kong, A., G. Thorleifsson, H. Stefansson, G. Masson, A. Helgason, etal., 2008 Sequence variants in the RNF212 gene associate with genome-wide recombination rate. Science 319: 1398-1401.

35. Li, X.-M., C. Yu, Z.-W. Wang, Y.-L. Zhang, X.-M. Liu, et al., 2013 DNA Topoisomerase II Is Dispensable for Oocyte Meiotic Resumption but Is Essential for Meiotic Chromosome Condensation and Separation in Mice. Biol. Reprod. 89: 118, 1-11.

36. Ma, L., J. R. O'Connell, P. M. VanRaden, B. Shen, A. Padhi, et al., 2015 Cattle Sex-Specific Recombination and Genetic Control from a Large Pedigree Analysis. PLoS Genet. 11:e1005387.

37. Moskvina, V., and K. M. Schmidt, 2008 On multiple-testing correction in genome-wide association studies. Genet. Epidemiol. 32: 567-573.

38. Muller, H., 1964 The relation of recombination to mutational advance. Mutat. Res. 1: 2-9.

39. Munoz-Fuentes, V., M. Marcet-Ortega, G. Alkorta-Aranburu, C. Linde Forsberg, J. M. Morrell, et al., 2015 Strong Artificial Selection in Domestic Mammals Did Not Result in an Increased Recombination Rate. Mol. Biol. Evol. 32: 510-523.

40. Myers, S., L. Bottolo, C. Freeman, G. McVean, and P. Donnelly, 2005 A fine-scale map of recombination rates and hotspots across the human genome. Science 310: 321-4.

41. Nagamine, y., R. Pong-Wong, P. Navarro, V. Vitart, C. Hayward, et al., 2012 Localising Lociunderlying Complex Trait Variation Using Regional Genomic Relationship Mapping. PLoS ONE 7: e46501.

42. O'Connell, J., D. Gurdasani, O. Delaneau, N. Pirastu, S. Ulivi, et al., 2014 A General Approach for Haplotype Phasing across the Full Spectrum of Relatedness. PLoS Genet. 10: e1004234.

43. Otto, S. P., and N.H. Barton, 2001 Selection for recombination in small populations. Evolution 55:1921-1931.

44. Otto, S. P., and T. Lenormand, 2002 Resolving the paradox of sex and recombination. Nat. Rev. Genet. 3: 252-261.

45. Parisi, S., M. J. McKay, M. Molnar, M. A. Thompson, P. J. van der Spek, et al., 1999 Rec8p, a Meiotic Recombination and Sister Chromatid Cohesion Phosphoprotein of the Rad21p Family Conserved from Fission Yeast to Humans. Mol. Cell. Biol. 19: 3515-3528.

46. Petit, M., J.-M. Astruc, J. Sarry, L. Drouilhet, S. Fabre, et al., 2017 Variation in Recombination Rate and Its Genetic Determinism in Sheep Populations. Genetics 207: 767-784.

47. Qiao, H., H. B. D. P. Rao, Y. Yang, J. H. Fong, J. M. Cloutier, et al., 2014 Antagonistic roles of ubiquitin ligase HEI10 and SUMO ligase RNF212 regulate meiotic recombination. Nat. Genet. 46: 194-199.

48. Reinink, P., and I. Van Rhijn, 2009 The bovine T cell receptor alpha/delta locus contains over 400 V genes and encodes V genes without CDR2. Immunogenetics 61: 541-549.

49. Reynolds, A., H. Qiao, Y. Yang, J. K. Chen, N. Jackson, etal., 2013 RNF212 is a dosagesensitive regulator of crossing-over during mammalian meiosis. Nat. Genet. 45: 269-78.

50. Rónnegàrd, L., S. E. McFarlane, A. Husby, t. Kawakami, H. Ellegren, etal., 2016 Increasing the power of genome wide association studies in natural populations using repeated measures: evaluation and implementation. Methods Ecol. Evol. 7: 792-799.

51. Sandor, C., W. Li, W. Coppieters, t. Druet, C. Charlier, etal., 2012 Genetic variants in REC8, RNF212, and PRDM9 influence male recombination in cattle. PLoS Genet. 8: 1-13.

52. Shin, J.-H., S. Blay, B. McNeney, and J. Graham, 2006 Ldheatmap: An r function for graphical display of pairwise linkage disequilibria between single nucleotide polymorphisms. J. Stat. Soft. 16: Code Snippet 3.

53. Stapley, J., P. G. D. Feulner, S. E. Johnston, A. W. Santure, and C. M. Smadja, 2017 Variation in recombination frequency and distribution across Eukaryotes: patterns and processes. Philos. Trans. R. Soc. Lond., B, Biol. Sci. 372: 20160455.

54. Theodosiou, L., W. O. McMillan, and O. Puebla, 2016 Recombination in the eggs and sperm in a simultaneously hermaphroditic vertebrate. Philos. Trans. R. Soc. Lond., B, Biol. Sci. 283: 20161821.

55. Wang, Z., B. Shen, J. Jiang, J. Li, and L. Ma, 2016 Effect of sex, age and genetics on crossover interference in cattle. Sci. Rep. 6: 37698.

56. Yang, J., S. H. Lee, M. E. Goddard, and P. M. Visscher, 2011 GCTA: a tool for genome-wide complex trait analysis. Am. J. Hum. Genet. 88: 76-82.

57. Yang, S.-W., C. Gao, L. Chen, Y.-L. Song, J.-L. Zhu, etal., 2012 NEK9 regulates spindle organization and cell cycle progression during mouse oocyte meiosis and its location in early embryo mitosis. Cell Cycle 11: 4366-4377.

58. Zerbino, D. R., P. Achuthan, W. Akanni, M. Amode, D. Barrell, etal., 2018 Ensembl 2018. Nucleic Acids Res. 46: D754-D761.

